# Phylogenomic assessment of microhylid frogs reveals widespread taxonomic confusion in the Asterophryinae and establishes the timing of diversification in Australia

**DOI:** 10.64898/2026.01.05.697806

**Authors:** Ian G. Brennan, Conrad J. Hoskin, Stephen J. Richards, Alan R. Lemmon, Emily Moriarty Lemmon, Stephen C. Donnellan, J. Scott Keogh

**Affiliations:** Division of Ecology & Evolution, Research School of Biology, Australian National University, Canberra, ACT 2601, Australia; Natural Sciences, Queensland Museum, Brisbane, QLD 4011, Australia; College of Science and Engineering, James Cook University, Townsville, QLD 4811, Australia; South Australian Museum, North Terrace, Adelaide, SA 5000, Australia; Department of Biological Science, Florida State University, Tallahassee FL 32306, USA; Australian Museum Research Institute, Australian Museum, 1 William Street, Sydney. NSW 2010, Australia

**Keywords:** amphibia, Australasia, narrow-mouthed frogs

## Abstract

Microhylid frogs are a hyper-diverse family thought to have radiated explosively around the Cretaceous-Paleogene boundary. Roughly half of microhylid species richness is concentrated into a single subfamily, the Asterophryinae, which is centered in New Guinea and surrounds, and has been a rich source for species discovery over the past 50 years. However, resolving Asterophryinae phylogenetics has remained a challenge, with frequent taxonomic reshuffling. To address this instability, we generated a sequence-capture molecular dataset to investigate the phylogenetics of the group. This included 71 species of Asterophryinae, across 13 of 17 recognized genera representing extensive sampling of the New Guinea radiation and full sampling of Australian microhylid species. Our dated species tree supports an explosive diversification of microhylids in New Guinea near the start of the Miocene, approximately 20 million years ago. Asterophryinae expansion into northern Australia occurred much later (∼10 ma) and is marked by well supported clades of *Austrochaperina* and *Cophixalus* that show temporally consistent splits from their New Guinea sister taxa. Our phylogeny allows us to identify several instances of polyphyly, which are at odds with our current understanding of intergeneric relationships within the Asterophryinae. We suggest that this confusion is a result of rapid radiation and morphological variability across some poorly defined genera. This work establishes a reliable phylogenetic framework that can form a foundation for a more stable taxonomy of the Asterophryinae.

**Highlights:**

- A phylogenomic assessment of the globally distributed frog family Microhylidae
- The subfamily Asterophryinae radiated explosively in New Guinea
- Australian species represent two distinct clades
- Assignments of species to genera by morphological means are often unreliable

## Introduction

Narrow-mouthed frogs, Family Microhylidae, are one of the most species-rich groups of amphibians worldwide. These tropical and subtropical frogs comprise nearly 800 species with a wide array of ecologies and morphologies (Frost, 2026). Their diversity is so broad that it defies easy summary but includes everything from round-bodied fossorial to slender toe-padded arboreal species, desert and rainforest specialists, and a variety of larval strategies (aquatic, direct-developing, foam-nest living, non-feeding). They are also geographically widespread, occurring from North, Central and South America, Africa, Madagascar, India, and southeast Asia, to New Guinea and northern Australia (Fig.1). Microhylids are allocated to 12 subfamilies, and the relationships within and between them have been the subject of many molecular phylogenetic assessments (van der Meijden et al. 2007; Kurabayashi et al. 2011; da Sá et al. 2012; Peloso et al. 2016; Tu et al. 2018; Streicher et al. 2020; Portik et al. 2023a; Portik et al. 2023b) (see Peloso et al. 2016 Fig.1). The increasing size of molecular datasets has revealed support for recognized subfamilies and provided evidence for some intrafamilial relationships, despite using different marker types (Feng et al. 2017; Streicher et al. 2020; Hime et al. 2021). This level of agreement, however, has not extended to relationships within the most species-rich subfamily—the Asterophryinae.

**Figure 1.**
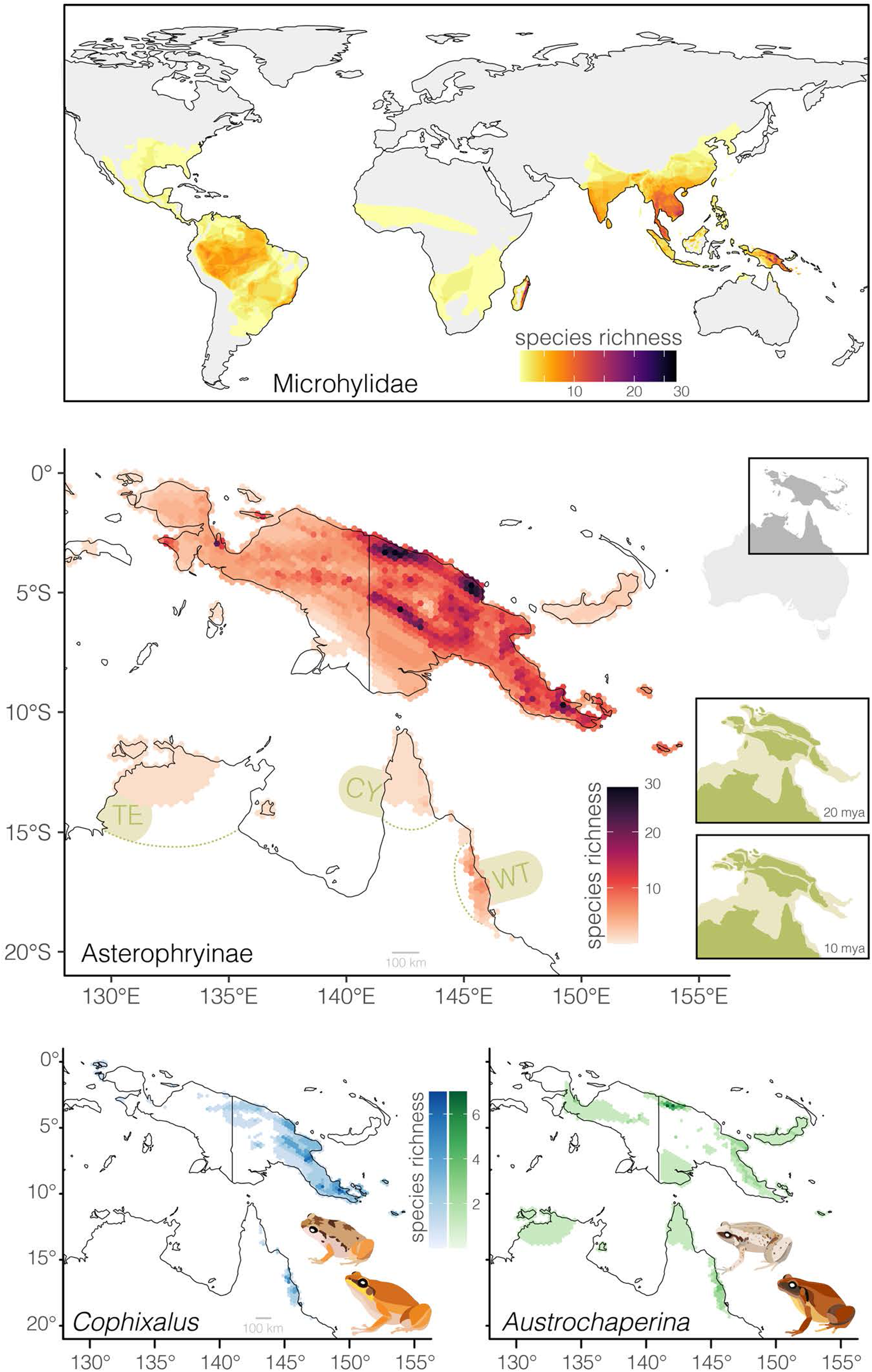
The Microhylidae are a global frog radiation with multiple hotspots of species richness. (Top) Mapping the global distribution of microhylid species highlights high richness on the east coast of Madagascar, across southeast Asia, and particularly on New Guinea. (Middle) Across New Guinea, richness is dominated by the subfamily Asterophryinae, and surrounding islands and landmasses show comparatively low diversity. Australian regions mentioned in the text are delineated by dotted lines: Top End (TE), Cape York (CY), Wet Tropics (WT). Inset maps at right indicate the proximity and connectivity of Australia and the various geological blocks that ultimately formed New Guinea at two different time periods: 20 million years ago during the rapid diversification of Asterophryinae microhylids in this region, and 10 million years ago when *Austrochaperina* and *Cophixalus* are hypothesized to have dispersed onto the Australian continent. Geological reconstructions are derived from Gold et al. 2020. (Bottom) The contemporary distribution of microhylids in Australia is concentrated in the Wet Tropics (WT) of North Queensland. Maps at bottom indicate the spatial dynamics of species richness in *Cophixalus* (left) and *Austrochaperina* (right). One Australian species *Austrochaperina gracilipes* is also found in southern New Guinea.

The Asterophryinae is the largest subfamily of microhylids comprising over 370 species and 17 genera. Asterophryines are distributed from mainland southeast Asia (*Siamophryne*, *Vietnamophryne*), to Borneo (*Gastrophrynoides*), the Philippines (*Aphantophryne*), New Guinea (14 genera), and northern Australia (*Austrochaperina, Cophixalus*) (Frost, 2026). Richness peaks in New Guinea (Fig.1), with over 250 species and likely many more to be described (Ferreira et al. 2024; Ferreira et al. 2025). In contrast, microhylid diversity in Australia is limited to ∼25 species, with just two genera (*Austrochaperina* and *Cophixalus*) on the mainland (Fig.1; Zweifel 1985; Hoskin 2004; Hoskin 2013), and two recently described species (*Callulops gobakula* and *Choerophryne koeypad*) endemic to Dauan Island, politically part of Australia, but geographically adjacent to New Guinea (Hoskin 2025). The vast majority of microhylid diversity in Australia comprises *Cophixalus* species in the Wet Tropics rainforests, with smaller numbers of species in the drier Cape York region and a single species of *Austrochaperina* in the far north of the Northern Territory (Fig. 1). The evolutionary history of microhylid frogs in Australia is currently unresolved, including whether mainland Australian species are the result of repeated dispersals from New Guinea, or instead represent Australian clades.

Phylogenetic relationships within and among genera of the Asterophryinae have been examined thoroughly, but with little consensus, hindered by frequent findings of para- and polyphyly (Zweifel, 1972; Kohler and Gunther, 2008; Rivera et al. 2017; Hill et al. 2022, Hill et al. 2023). This is likely due to rapid radiation of the group after crossing Wallace’s Line ∼20 million years ago, resulting in bursts in speciation and ecomorphological diversification. As a result of highly variable morphologies, and because it is rare for diagnoses of Asterophryinae species to include molecular evidence, generic assignments in this subfamily have changed frequently (Frost, 2026). The frustrations of Asterophryinae generic assignments have been so extreme that Dubois et al. (2021) proposed that all 360+ species (with exception of *Gastrophrynoides, Siamophryne,* and *Vietnamophryne*) be lumped under a single genus, *Asterophrys*. This suggestion, however, has not been adopted by researchers in the field (Frost, 2026).

Here we present a phylogenomic perspective on the diversification of Asterophryinae microhylids, particularly among New Guinean and Australian taxa. We started by generating a sequence-capture dataset to investigate the topology and timing of Sahulian asterophryine diversification, with the goal of providing a reliable backbone of intergeneric relationships. We assess the timing of diversification in New Guinea and Australia, and we resolve how the Australian species fit into the evolutionary history of this group. While we know a great deal about many aspects of Australian frog biology (Tyler 1998; Anstis 2017, Brennan et al. 2023), comparatively little is known about the phylogenetics and history of the ∼25 species of microhylids (Zweifel 1985; Hoskin 2004). We also evaluated commonly used morphological information to quantify the distribution and utility of traits in generic assignments. Our work aims to provide insight into the phylogenetics of a taxonomically volatile group and expand our understanding of the biotic interchange between Australia and New Guinea.

## Materials and Methods

### Visualising Species Richness

To summarize geographic patterns of species richness in microhylid frogs we started by downloading species range maps for all Anurans from the IUCN Red List spatial data website (2026). We subset these data separately to the family Microhylidae, Asterophryinae genera, and the genera *Austrochaperina* and *Cophixalus*. We summed richness at a resolution of one quarter degree and visualised the resulting raster using *ggplot2* (Wickham 2011). All code is available in the supplement.

### Phylogenomics

We assembled a sequence-capture dataset comprising 149 frog samples across 107 species that span nearly all microhylid subfamilies (10 of 12 recognized). Exceptions are limited to Hoplophryninae and Melanobatrachinae frogs of East Africa and India. Sampling focused on the Asterophryinae and represents 71 species from 13 of 17 recognized genera (with exceptions *Gastrophrynoides*, *Paedophryne*, *Siamophryne*, *Vietnamophryne*) (Table S1). We include near-complete sampling of the Australian Asterophryinae species (5 *Austrochaperina* spp.; 19 *Cophixalus* spp.) with the exception of *Cophixalus peninsularis*, which is known only from two specimens collected in the 1980s and is likely to be conspecific to *C. crepitans* (Hoskin, 2004).

We generated new Anchored Hybrid Enrichment (AHE—Lemmon et al. 2012) data for 96 samples and combined these with outgroup samples from Hime et al.’s (2021) amphibian phylogenomic dataset. We initiated this process by blasting AHE loci against the *Xenopus tropicalis* genome using *metablastr* (*blast_best_reciprocal_hit*) (Benoit & Drost 2021) and renaming loci according to their orthologs in *Xenopus*. We similarly carried out this process on anuran samples from Hime et al. (2021) to harmonize target sequences across datasets. Samples across different AHE projects were combined using the *pipesnake* workflow (Brennan et al. 2024) to align and trim sequence data, and estimate locus and species trees (using *--stage from-prg*). Briefly, sequences were aligned with *mafft* (Katoh et al. 2013), trimmed for gappy sites using *clipkit* (Steenwyk et al. 2020), then locus trees (n=450) were estimated under maximum-likelihood in IQTREE2 (Minh et al. 2020). Best fitting models of nucleotide substitution were identified by ModelFinder (Kalyaanamoorthy et al. 2017), and IQTREE2 fit 1,000 ultrafast bootstraps (Minh et al. 2013), before being passed to hybrid weighted-ASTRAL (Zhang et al. 2018) to estimate a species tree.

To verify the identity of newly sequenced samples we assembled off-target reads of the mitochondrial loci *CYTB* and *ND4* and combined these data with the alignments of Hill et al. (2023). We started by loosely mapping raw sequence reads to the *Microhyla pulchra* mitochondrial genome using *BBMAP* (Bushnell, 2014), then assembled mapped reads using *SPAdes* (Prjibelski et al. 2020). We added new sequences to the existing alignments with *mafft*, concatenated the alignments, and estimated a single mitochondrial topology using IQTREE2.

### Divergence Dating

To estimate divergence times among taxa on the ASTRAL species tree we applied a series of fossil calibrations first compiled by Feng et al. (2017) (Table S2), and used the Bayesian divergence time software MCMCtree (Rannala & Young 2007). We started by downloading orthologous coding sequences from the Orthologous Matrix (Altenhoff et al. 2024) for *Xenopus* and *Bufo* and aligned the exonic loci via MACSE (Ranwez et al. 2018). We then added our ingroup microhylid sequences to these codon-aligned sequences via *mafft* (*--add*, *--keeplength*), concatenated them, and partitioned first and second codon positions together following the strategy of dos Reis et al. (2018). Complex partitioning strategies such as filtering by evolutionary rate are possible but less influential than the absolute number of partitions (dos Reis et al. 2012). Additional data partitions ultimately incur substantial computational costs for modest increases in dating precision, and so we opted instead for a more conservative approach. We then used *baseml* to estimate approximate likelihoods (dos Reis & Yang 2011) and branch lengths before running *mcmctree* on the gradient and Hessian (in.BV file) for four replicate analyses. We inspected mcmc files for stationarity and compared for convergence, then combined them using logCombiner, and used this combined mcmc file to summarize divergence times on our tree (*print = -1* in .ctl file). Sample, alignment, and gene trees are available alongside all other materials on Dryad (doi:[TO_BE_UPDATED]) and GitHub (https://github.com/IanGBrennan/Asterophryinae).

### Taxonomy

For consistency, we adopt the taxonomy of Amphibian Species of the World v6.2 (Frost, 2026), which shows minor differences in the number of recognized species, recognized genera, and generic assignments from other amphibian authorities such as AmphibiaWeb. Notably, Amphibian Species of the World has incorporated the taxonomic suggestions of Rivera et al. (2017) regarding assignment of Asterophryinae genera and has not implemented the recommendation of Dubois et al. (2021) to lump most asterophryines into *Asterophrys*.

### Morphology

We quantified commonly collected morphological traits used in Asterophryinae species descriptions, to evaluate a potential explanation for why generic assignments have been so unstable. Asterophryinae genera diagnoses often rely on both external linear measurements and internal anatomical traits such as presence/absence of clavicles, procoracoids, musculature, and the shape of the jaw. However, internal traits are not always assessed for new species because this practice is time consuming and destructive. We started by collecting eight morphological measurements from species descriptions in the microhylid literature (40 publications, see *Supplement*): snout-urostyle length (SUL), tibia length (TL), head width (HW), internasal distance (IN), eye-nasal distance (EN), eye diameter (EYE), third finger disc diameter (F3D), and fourth toe disc diameter (T4D). To remove the effect of size on individual traits (allometry) we calculated the geometric mean of all traits by individual and used this to transform measurements into log-shape ratios. We retained the geometric mean as a ninth trait (SIZE). We then used current generic assignments as a discrete character and carried out a RandomForest (Liaw & Wiener, 2022) analysis across 10,000 decision trees on the morphological traits to determine rates of mischaracterization based on gross phenotype (genus ∼ morphology). For comparison with other commonly used methods, we also applied Linear Discriminant (LDA) and Flexible Discriminant (FDA) analyses. To visualize the partitioning of morphological space we used dimensionality reduction techniques (PCA, LDA, FDA) and plotted the first two axes of variation, which together account for ∼89%.

## Results

### Phylogenomics and Divergence Dating

We present a phylogenetic hypothesis of relationships for 107 species of microhylid frogs based on new anchored hybrid enrichment (AHE) data for 95 new samples. Sampling includes most recognized Asterophryinae genera (13 of 17) and, for the first time, all mainland Australian species of Asterophryinae. We combine this new molecular dataset with existing AHE data from Hime et al. (2021) to estimate relationships among 10 of the 12 microhylid subfamilies (Fig. 2; S1).

**Figure 2.**
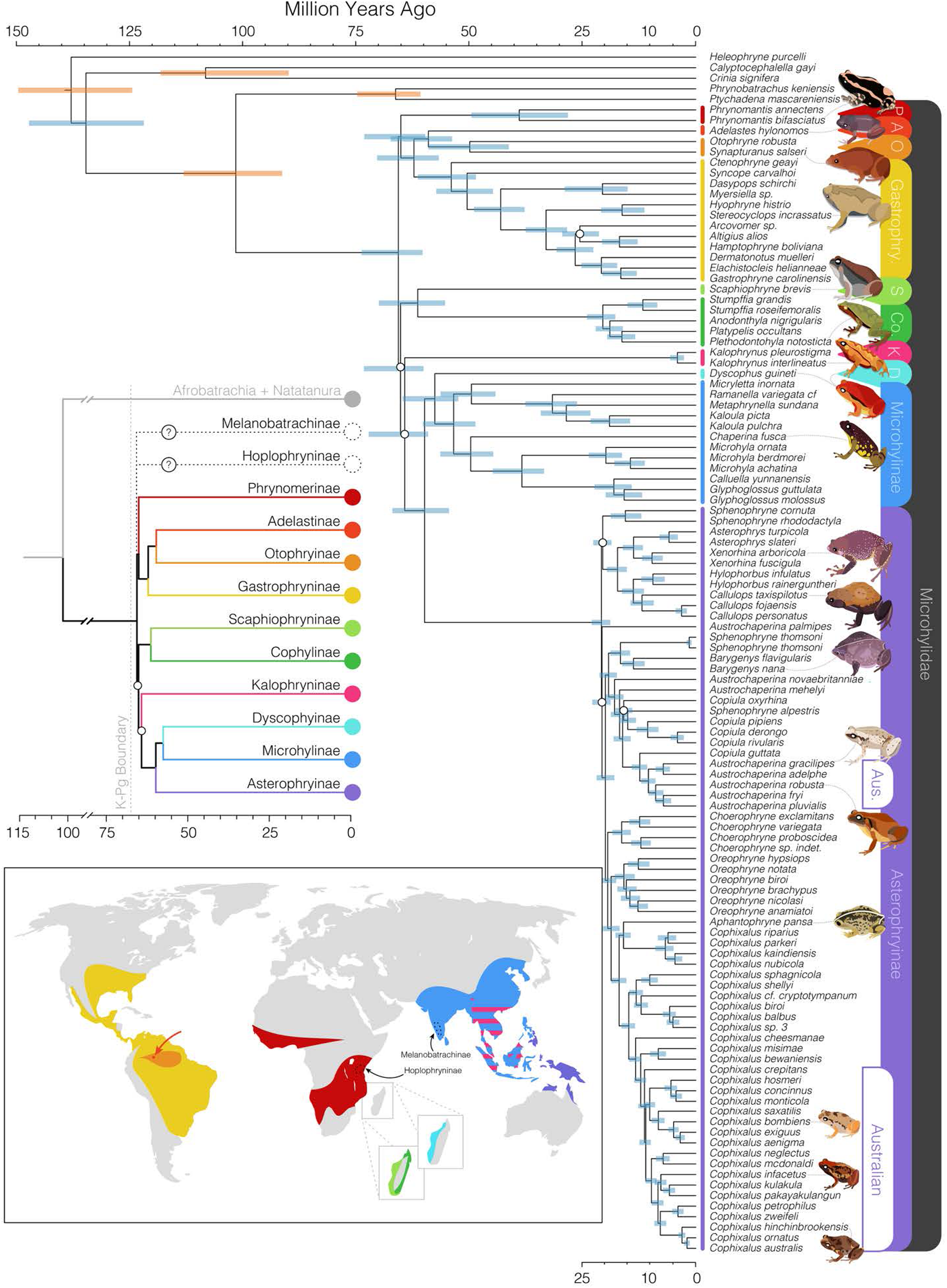
Time calibrated species tree of microhylid frogs highlights the global radiation of the family around the K-Pg boundary (∼66 mya) and subsequent explosive diversification of the subfamily Asterophryinae in the early Miocene (from ∼20 mya). Inset tree shows the relationships and divergence times among microhylid subfamilies, and color codings match to the geographic distributions of those clades on the map below. Note (1) the small distribution of Adelastinae indicated by an arrow in northern South America, (2) the overlapping distribution of Microhylinae and Kalophryninae in southeast Asia indicated by pink stripes, (3) the presence of three subfamilies on Madagascar, and (4) the uncertain phylogenetic position of the Hoplophryninae (east African) and Melanobatrachinae (south Indian). Primary tree at right shows species level relationships and divergence times, unlabelled nodes have posterior probabilities >0.9 in the weighted-ASTRAL species tree, white circles at nodes indicate branches with local posterior probabilities <0.9. The 95% Confidence intervals of divergence times are shown as shaded rectangles at nodes, with orange CIs indicating calibrated nodes.

We captured 356 loci with a mean alignment length of 1564 bp (max. = 5535, min. = 377) and sample occupancy of 120 individuals (max. = 148, min. = 34). Species tree analyses with ASTRAL provided strong support (>90 local posterior probability) for most branches of the tree (Fig. 2; S1), with some exceptions among subfamilial splits (e.g. position of Kalophryninae and Scaphiophryninae/Cophylinae relative to the South American and Asian clades) and among some Asterophryinae clades. The subfamily topology is entirely consistent with Hime et al. (2021) and differs only in a small number of changes from Feng et al. (2017) (placement of Phrynomerinae) and Streicher et al. (2020) (position of Kalophryninae and Phrynomerinae). The microhylid stem branch is very long (>35 ma) and leads to an explosive radiation into major microhylid clades (subfamilies) roughly coincident with the Cretaceous-Paleogene boundary. The four earliest splits in the family all occur within a 1.5 million year window at this time (Fig. 2; S2).

Our topology for the Asterophryinae differs considerably from recent investigations by Rivera et al. (2017) and Hill et al. (2022), and in some ways bears greater similarity to Tu et al. (2018). To verify the identity of new species we successfully recovered off-target mitochondrial data for 84 of 95 new samples (Fig. S3). While our species sampling (∼70 spp.) represents roughly a third of the species included in those works, there are notable differences in relationships among major clades, which we will focus on. Importantly, we estimate *Oreophryne* is the deeply nested sister taxon to *Cophixalus*, and not an early branch of the tree as seen in Hill et al. (2022) (‘*Oreophryne* A’). We group *Aphantophryne pansa* with *Cophixalus* and not *Oreophryne* (‘*Oreophryne* B*’*). We do not find *Barygenys* embedded within *Austrochaperina*; rather, we identify a sister-group relationship between *Barygenys* and *Sphenophryne (Genyophryne) thomsoni*. Some topological differences cannot be addressed with our smaller taxon sampling, such as the monophyly of *Oreophryne.* There are, however, important similarities, which lend support to an emerging consensus in the intergeneric relationships of Asterophryinae. *Asterophrys*, *Callulops*, *Hylophorbus*, and *Xenorhina* are consistently recovered as a clade. While not included in this work, there is consistent support from other datasets that *Mantophryne* also sits in this clade (Morris et al. 2026). *Austrochaperina* is clearly paraphyletic, appearing as four lineages/clades in our tree. *Austrochaperina palmipes* is a highly divergent lineage not closely related to other *Austrochaperina* or any other single group. *Sphenophryne* is also clearly paraphyletic, appearing in three positions in the tree. *Cophixalus* (here including *Aphantophryne pansa*) forms a well supported New Guinean and Australian clade.

The timing of Asterophryinae diversification is rapid and resembles the rapid pace of diversification among microhylid subfamilies. A long stem branch that spans ∼40 ma was followed by explosive diversification around 20 million years ago and was quickly followed by eight early splits which occurred within a 1.5 million year window. Asterophryinae expansion into northern Australia occurred much later (∼10 mya) and is marked by well supported clades of *Austrochaperina* and *Cophixalus* that show contemporaneous splits from their New Guinea sister lineages. The internal topologies of these clades are broadly consistent with their last phylogenetic assessment two decades ago (Hoskin 2004).

Our morphological dataset covered ∼532 individual asterophryine frogs representing 143 species (∼30% of species diversity) and all 17 recognized genera. Visualizations of dimensionality-reduced morphological data (PCA, LDA, FDA) identified large regions of morphospace shared among genera, with some highly distinct forms (*Barygenys*, *Paedophryne*, *Xenorhina*) (Fig. 3). Some groups (*Choerophryne*, *Cophixalus*, *Sphenophryne*) show large or discontinuous distributions that indicate variable morphologies (Fig. 3). Classification error for generic assignments using RandomForests were bimodal, though typically low (<10%) (Fig. S4). However, five genera (*Copiula, Choerophryne, Oreophryne, Mantophryne, Siamophryne*) show moderate (10<x<30%) rates of error and three others (*Aphantophryne, Sphenophryne, Vietnamophryne)* show high rates (30–100%). This suggests that morphological characters often used to describe genera do not differentiate them cleanly, even in combination.

**Figure 3.**
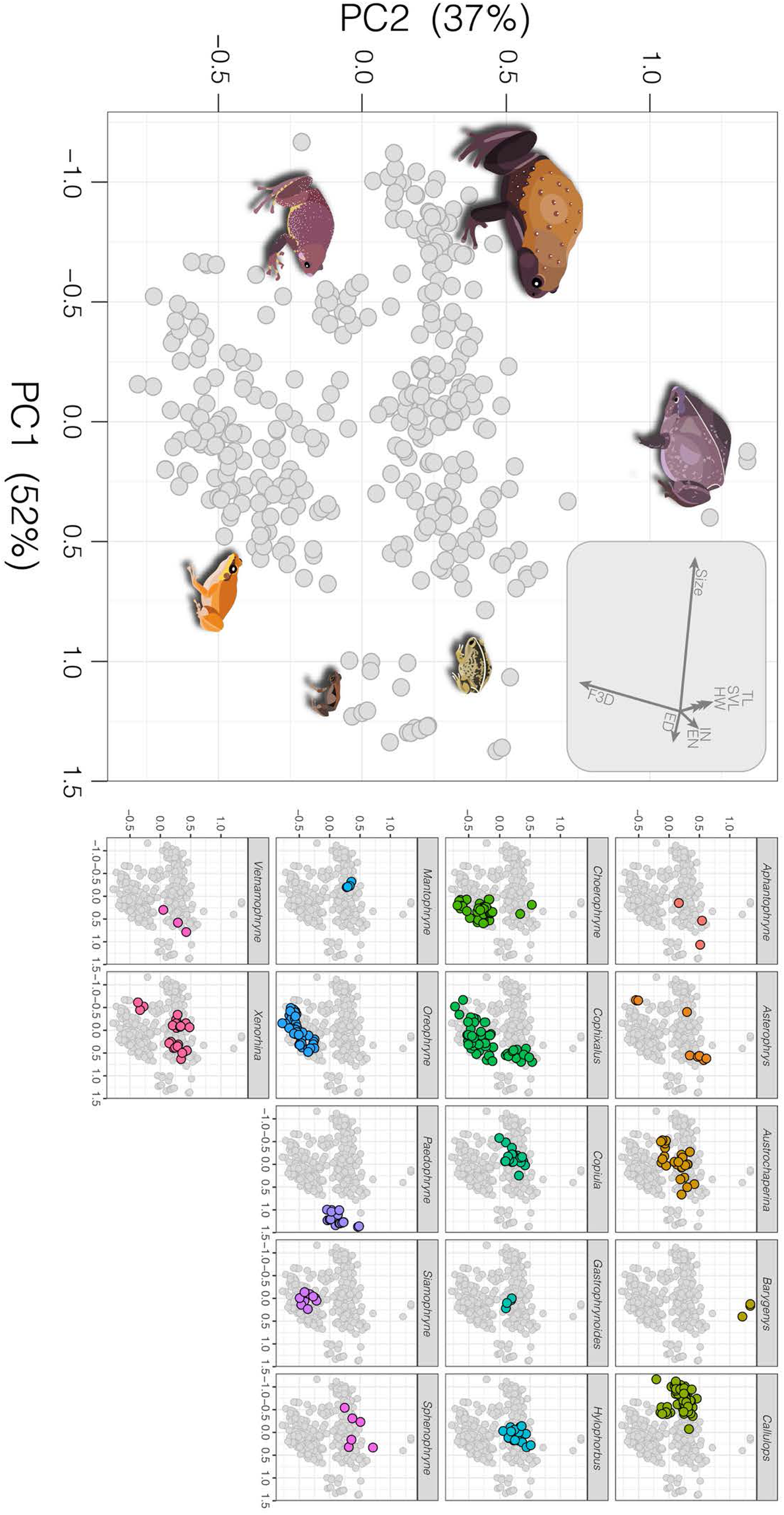
Asterophryinae morphospace shows considerable overlap among genera, with a small number of highly divergent groups (*Barygenys*, *Callulops*, *Paedophryne*). This morphospace, approximated by linear measurements, does not allow for a neat assignment of species to genera through traditional discriminant means (machine learning, linear or flexible discrimination). We suggest that this result underlies why some genera remain difficult to diagnose and species assignments are unstable. (Left) PCA of first two principal components which together explain ∼90% of the variation. Diversity is primarily described by size and the diameter of the toe pad on the third finger, but there are obvious other morphological axes of diversity as seen by the species silhouettes. (Right) the same PCA, but highlighting the distribution of genera within this space.

## Discussion

Microhylids comprise the third-largest family of living amphibians, with almost 800 species distributed across the tropics. Phylogenetics of this group have been a popular topic over the last two decades, greatly improving our understanding of the patterns of diversification and taxonomy of the group (van der Meijden et al. 2007; Kurabayashi et al. 2011; da Sá et al. 2012; Peloso et al. 2016; Tu et al. 2018; Streicher et al. 2020). However, much still remains to be discovered, resolved, and described (Ferreira et al. 2025). Recent global amphibian phylogenomics initiatives (Feng et al. 2016; Hime et al. 2020) have helped to establish the evolutionary context in which microhylids have diversified, and have built substantial data sets to test earlier hypotheses that were based on fewer loci. Our microhylid phylogenomics study presented here helps to clarify our understanding of this globally distributed frog group and provides substantial new information on the diverse Asterophryinae, especially in New Guinea and Australia.

Phylogenetic relationships and divergence times among microhylid subfamilies estimated here are largely consistent with other phylogenomic investigations (Feng et al. 2017; Streicher et al. 2020; Hime et al. 2021; Portik et al. 2023a). We confirm an explosive radiation at the base of the microhylid tree, rapidly separating the subfamilies. This event is coincident with, or closely follows, the K-Pg turnover. The timing of microhylid crown diversification is important because its ancestral lineage split from the rest of Ranoidea nearly 100 million years ago. In light of this, the long stem branch leading to the Microhylidae (>35 ma) is likely indicative of elevated Cretaceous extinction and a dramatic rebound in the Paleogene. This pattern is seen in many vertebrate groups, such as squamate reptiles (Longrich et al. 2012), mammals (Meredith et al. 2011), and frogs (Feng et al. 2017), and potentially highlights biased survival of nocturnal groups (Wu et al. 2017). Importantly, the rapid diversification of the group in the wake of the K-Pg turnover—including multiple splits in the first 1.5 ma and establishment of all subfamilies in <10 ma—has the unfortunate effect of blurring the true branching order of the tree. One consequence is ambiguity in the position of Kalophryninae and Cophylinae/Scaphiophryninae. Frequent successive speciation events can be difficult for molecular phylogenetic methods to resolve because high levels of incomplete lineage sorting and gene tree incongruence can obscure the topology (Linkem et al. 2016).

Our topology and divergence dating offers an opportunity to assess the biogeographic history of an old, diverse group, with a wide distribution. Looking across our tree, we see among-subfamily relationships that might best reflect a Gondwanan history. For example, North and South American subfamilies (Adelastinae, Otophryninae, Gastrophryninae) form the sister taxon to an African group (Phrynomerinae), and Madagascan and East Asian clades are sister taxa. However, the timing of microhylid diversification is at odds with a Gondwanan diversification-by-vicariance scenario. The rapid splitting of subfamilies from one another occurred around the Cretaceous-Paleogene boundary roughly 66 million years ago, long after the separation of Africa, Madagascar and India from Gondwana, ruling out Gondwanan vicariance as the primary driver of contemporary distributional patterns. Instead, the history of this group has likely been driven by significant long-distance dispersal events followed by subsequent diversification. The number and nature of these events seem almost implausible. For example, van der Meijden et al. (2007) suggested that better taxon sampling might resolve the relationships among the Microhylinae (Southeast Asia), Asterophryinae (eastern Malesia & New Guinea), and Dyscophinae (Madagascar). We interpret their comment to suggest that increased phylogenetic resolution might cluster the Microhylinae and Asterophryinae and provide a more parsimonious biogeographic story. Instead, more and better data presented here confirm the topology that they first presented (Asterophryinae as the sister taxon to Dyscophinae and Microhylinae), emphasizing the convoluted dispersal history of this group. As a result, microhylids either undertook two separate dispersals from Madagascar to Asia and Australia, or a single dispersal followed by a return of the Dyscophinae to Madagascar, a trip of ∼4,000 km. Long distance dispersals are not unheard of in amphibians (Fonte et al. 2019), but these distances over open ocean would be exceptional. Like van der Meijden et al. nearly two decades ago, we also find ourselves wishing for more molecular data and more comprehensive taxon sampling to pair with powerful contemporary biogeographic models that would “unambiguously resolve the biogeographic history of the Microhylidae.”

Among the Microhylidae, nearly half of all species richness belongs in the Asterophryinae. The majority of this richness is concentrated in and around New Guinea; however, there is also an early branching clade (not included in our study) spread across Vietnam and Thailand (*Siamophryne, Vietnamophryne*), peninsular Malaysia (*Gastrophrynoides*), and on Borneo (*Gastrophrynoides*) (Kurabayashi et al. 2011; Suwannapoom et al. 2018; Poyarkov et al. 2018; Portik et al. 2023a). Given that the closest relatives of the Asterophryinae are likely the Microhylinae of Asia (and inexplicably the Dyscophinae of Madagascar), it seems plausible that asterophryines dispersed via a stepping stone procession from southeast Asia, crossing Wallace’s Line just once (see Fig. 1 of Poyarkov et al. 2018). Continued dispersal among island groups within Malesia and expansion across New Guinea have likely continued to drive species richness. As it currently stands, there are more than 250 described species found on New Guinea, across a diversity of habitats. This naturally has led to the idea that ecological opportunity has strongly contributed to asterophryine ecomorphological diversification (Rivera et al. 2017; Hill et al. 2022). Ecological and resulting morphological transitions are observable at deep timescales among genera (e.g. arboreal *Oreophryne,* fossorial *Xenorhina*) but also at shallow timescales (e.g., the arboreal *Xenorhina arboricola*, in an otherwise terrestrial genus *Xenorhina*). This process is ongoing, and species discovery rates in the Asterophryinae remain remarkably high (Ferreira et al. 2024; Ferreira et al. 2025). Here, we show that this incredible diversity has arisen within the last 20 million years.

The explosive diversification of New Guinean microhylids has resulted in relationships among genera that are poorly resolved, and further undermined by unreliable generic diagnoses (Zweifel 1972; Zweifel 1985; Burton 1986; Zweifel 2000). The vast majority of Asterophryinae species have been described by morphological investigation and await molecular assessment. Typically, species are assigned to genera based on suites of anatomical features, including osteological characters. However, these too may be more variable than anticipated. This is highlighted by the discovery by Günther et al. (2023) that two *Cophixalus* species possess procoracoids, where lack of these elements was previously considered diagnostic of the genus. Here we show (Fig. 3) that standard linear measurements—which contribute to generic and species diagnoses and commonly help to assign species to genera—often fail to accurately diagnose genera. This suggests that perhaps multiple forces are at play in causing the taxonomic confusion we see in Asterophryinae: (1) not all species have been thoroughly examined both externally and internally to fit them to an appropriate genus, (2) existing morphological diagnoses of genera are not reflective of our contemporary understanding of Asterophryinae diversity, and (3) rapid diversification may result in morphological patterns that do not match the underlying evolutionary tree (e.g. through hemiplasy). Regardless of what is causing this confusion, it is clear that some genera fill highly divergent and identifiable morphospaces, such as *Barygenys*, *Callulops*, and *Paedophryne*. Many others overlap considerably, complicating easy generic assignments. This is particularly the case for morphologically variable groups, such as *Cophixalus, Oreophryne*, and *Sphenophryne*. Others like *Austrochaperina* and *Copiula* are morphologically conservative but are not easily differentiated from one another. The result is a perfect storm where taxonomic uncertainties persist, muddled further by variable morphologies, and limited molecular phylogenetic information (sampling of both species and genetic loci).

While our data support prior studies in identifying a clade comprising *Asterophrys*, *Callulops*, *Hylophorbus*, *Mantophryne* (not sampled here), and *Xenorhina*, the number and position of remaining genera are hard to determine. Our phylogenomic perspective nests *Aphantophryne pansa* among *Cophixalus*, and together confidently associates them with *Choerophryne* and *Oreophryne* (but with limited species sampling in these two genera). Other assessments have identified more than one major clade of *Oreophryne* (Rivera et al. 2017; Hill et al. 2022; Portik et al. 2023a), however, our sampling lacks a representative of the *Oreophryne* ‘B’ clade. The assignment of species to *Austrochaperina*, *Copiula*, *Sphenophryne* and associated genera, however, remain inconsistent and clearly unresolved. While our topology is mostly well supported, it is important to note that our taxonomic sampling is incomplete, leaving a thorough taxonomic reassessment out of reach. While unsatisfying, it echoes previous calls for clarification of genus-level Asterophryinae taxonomy (Portik et al. 2023a). It is tempting to resolve the situation by lumping these genera all into *Asterophrys* following Dubois et al. (2021); however, we agree with Hill et al. (2022) and Frost (2026) in that this is not the preferred solution. A genus of that size (>375 spp.) and diversity would be unwieldy and fails to serve a purpose beyond being a monophyletic unit. Further phylogenomic sampling paired with a careful morphological assessment will be necessary to resolve outstanding Asterophryinae taxonomic issues.

The southwest Pacific is a region of complex geological and biogeographic history (Wallace, 1869; Hall, 1997). This area sits at the confluence of several continental plates that have been undergoing a complicated dance of movement and compression over millions of years. As a result, thousands of islands have emerged and subsided, including one of the world’s largest, New Guinea. The island of New Guinea itself is a composite formed by the amalgamation of several geologic terranes that accreted roughly around the same time as the explosive diversification of the Asterophryinae, ∼20 million years ago (Davies 2012; Gold et al. 2020). This suggests that the interplay of New Guinea’s foundational geologic units were influential in the radiation of asterophryine microhylids (Hill et al. 2023). We stress, however, that complex biogeographic scenarios, such as presented in Hill et al. (2023), must be confirmed in the presence of new phylogenetic evidence that may impact the evolutionary narrative.

Regardless of the order of movements among New Guinea’s biogeographic regions, it is evident that microhylids are capable dispersers, and our Australian-focused sampling adds a new layer to their history. We find evidence of two independent dispersals into Australia, one each for the genera *Austrochaperina* and *Cophixalus*. One taxon, *A. gracilipes*, occurs on both landmasses but sits in the Australian clade of species, suggesting a dispersal back to New Guinea (possibly during known land connections in the Pleistocene; Lewis et al. 2013). While the southern portion of New Guinea (the Australian Craton) has long been attached to the Australian continental plate, deeper time land connections (i.e., over millions of years) are poorly understood. Temporally consistent dispersal events from New Guinea to Australia by both *Austrochaperina* and *Cophixalus* suggest an elevated dispersal likelihood ∼10 million years ago. This was possibly the result of substantially lower sea levels that facilitated expansion not only into Cape York, but also the Top End of northern Australia (Fig. 1) (Miller et al. 2020). Our dating for microhylids is broadly consistent with dating of faunal interchange between other vertebrate groups between the two regions (e.g., marsupials, Mitchell et al. 2014). Once present in Australia, subsequent diversification of both *Austrochaperina* and *Cophixalus* has been centered in the Wet Tropics of Far North Queensland (Fig. 1). This wet and topographically complex landscape has long been a source of endemic diversification, likely a result of persistent rainforest habitats and sufficient refugia during periods of climatic volatility (Martin 2006; VanDerWal et al. 2009).

Our phylogenomic perspective on the Asterophryinae provides much needed resolution to many parts of the evolutionary tree, such as the timing and position of Australian species. Despite our progress, many questions remain for this species-rich and enigmatic group. Further molecular taxonomic sampling will undoubtedly help us to explore the complex biogeographic history of microhylids and unravel the amazing diversification of Asterophryinae on New Guinea.

## Data Statement

All data and code necessary to reproduce this work are available on GitHub at www.github.com/IanGBrennan/Asterophryinae

## Acknowledgments

We thank our many Australian partner institutions including museums and universities, and their associated curators and collections managers, who made this work possible through generous tissue loans and collections access. We wish to thank the technical staff at Florida State University for their support and hard work generating the genetic data presented here. J.S.K, C.J.H, and S.C.D thank the Australian Research Council for ongoing support. We appreciate provision of computing and data resources provided by the Australian BioCommons Leadership Share (ABLeS) program under the Australian Amphibian and Reptile Genomics initiative. These programs are co-funded by Bioplatforms Australia (enabled by NCRIS), the National Computational Infrastructure and Pawsey Supercomputing Centres. A sincere thank you to Editor-in-Chief Guillermo Ortí, Eli Greenbaum, and an anonymous reviewer for comments that improved the quality of this manuscript. We declare no conflicts of interest.

## Supplemental Information to

Document S1. Figures S1–SX and Tables S1–SX

## Supplementary Information

### Molecular Sampling

**Table S1:**
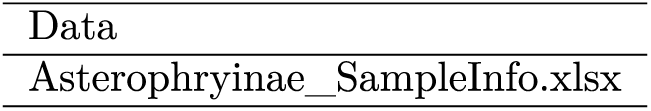
Table of molecular samples is included in ‘Sampling/Asterophryinae Sample-Info.xslx‘.

| Data |
| --- |
| Asterophryinae_SampleInfo.xlsx |

### Morphological Sampling

Below, we provide information about the collection of morphological traits from the litera-ture, as well as summary statistics about the sampling.

**Table S2:**
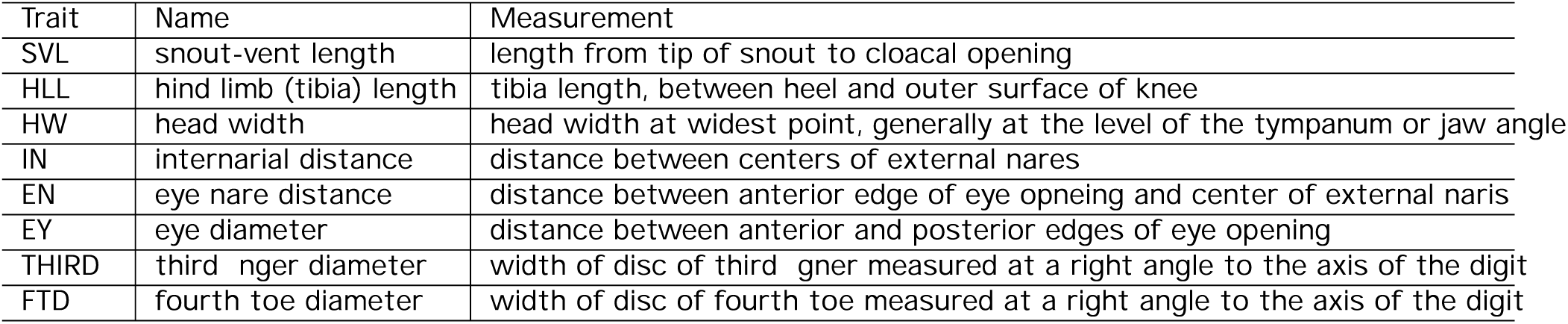
Morphological measurements of Asterophryinae frogs, following Zweifel (1972; 2000).

Morphological Classification

**Figure S1.**
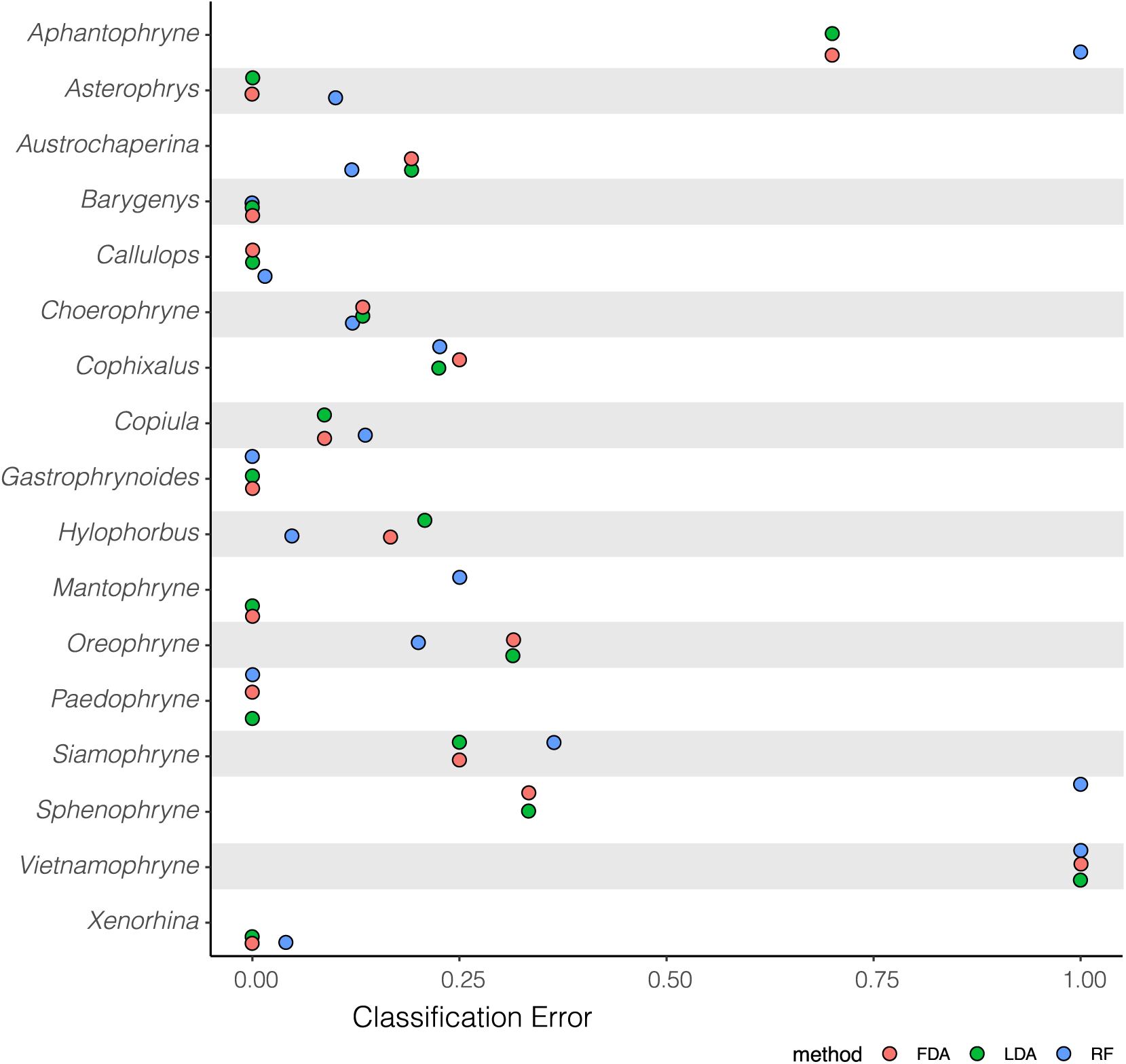
Classification error by Genus and classification method (color). Difference classi-fication methods (FDA–Flexible Discriminant Analysis; LDA–Linear Discriminant Analysis; RF–Random Forest) show comparable levels of classification error and are typically low, with the exception of some genera.

**Table S3:** Number of species (No.Species) and samples (No.Samples) for each genus included in the morphological analyses.

| Genus | No.Species | No.Samples |
| --- | --- | --- |
| Aphantophryne | 3 | 3 |
| Asterophrys | 4 | 11 |
| Austrochaperina | 16 | 27 |
| Barygenys | 3 | 4 |
| Callulops | 16 | 69 |
| Choerophryne | 7 | 34 |
| Cophixalus | 43 | 216 |
| Copiula | 7 | 26 |
| Gastrophrynoides | 2 | 4 |
| Hylophorbus | 10 | 22 |
| Mantophryne | 2 | 4 |
| Oreophryne | 9 | 48 |
| Paedophryne | 4 | 18 |
| Siamophryne | 1 | 11 |
| Sphenophryne | 5 | 6 |
| Vietnamophryne | 3 | 3 |
| Xenorhina | 8 | 26 |
| NA | 0 | 0 |

**Table S4:** Summary of pairwise model classifications across all three classification methods. Table is summarised as the actual genus of the sample (True Genus), the genus it was classified to (Assigned To), and the frequency with which individuals of that genus were classified to that specific other genus (Frequency), depending on method. Misclassifications are highlighted in red.

| True Genus | Assigned To | Frequency_RF | Frequency_LDA | Frequency_MDA |
| --- | --- | --- | --- | --- |
| Aphantophryne | Aphantophryne | 0 | 3 | 3 |
| Aphantophryne | Cophixalus | 2 | 5 | 5 |
| Aphantophryne | Paedophryne | 1 | 0 | 0 |
| Aphantophryne | Sphenophryne | 0 | 2 | 2 |
| Asterophrys | Asterophrys | 9 | 7 | 7 |
| Asterophrys | Callulops | 1 | 0 | 0 |
| Austrochaperina | Austrochaperina | 22 | 21 | 21 |
| Austrochaperina | Callulops | 0 | 1 | 1 |
| Austrochaperina | Choerophryne | 0 | 1 | 1 |
| Austrochaperina | Cophixalus | 2 | 0 | 0 |
| Austrochaperina | Copiula | 1 | 1 | 1 |
| Austrochaperina | Hylophorbus | 0 | 1 | 1 |
| Austrochaperina | Vietnamophryne | 0 | 1 | 1 |
| Barygenys | Barygenys | 3 | 3 | 3 |
| Callulops | Austrochaperina | 1 | 0 | 0 |
| Callulops | Callulops | 66 | 66 | 66 |
| Choerophryne | Choerophryne | 29 | 26 | 26 |
| Choerophryne | Cophixalus | 1 | 1 | 1 |
| Choerophryne | Hylophorbus | 1 | 0 | 0 |
| Choerophryne | Oreophryne | 2 | 3 | 3 |
| Cophixalus | Aphantophryne | 2 | 0 | 0 |
| Cophixalus | Asterophrys | 1 | 0 | 0 |
| Cophixalus | Austrochaperina | 1 | 2 | 2 |
| Cophixalus | Choerophryne | 1 | 0 | 0 |
| Cophixalus | Cophixalus | 41 | 31 | 30 |
| Cophixalus | Oreophryne | 5 | 5 | 5 |
| Cophixalus | Siamophryne | 2 | 2 | 2 |
| Cophixalus | Vietnamophryne | 0 | 1 | 1 |
| Copiula | Austrochaperina | 3 | 1 | 1 |
| Copiula | Copiula | 19 | 21 | 21 |
| Copiula | Hylophorbus | 0 | 1 | 0 |
| Copiula | Sphenophryne | 0 | 1 | 1 |
| Gastrophrynoides | Gastrophrynoides | 4 | 4 | 4 |
| Hylophorbus | Austrochaperina | 1 | 0 | 0 |
| Hylophorbus | Cophixalus | 0 | 2 | 2 |
| Hylophorbus | Hylophorbus | 20 | 19 | 20 |
| Hylophorbus | Sphenophryne | 0 | 1 | 1 |
| Hylophorbus | Vietnamophryne | 0 | 1 | 1 |
| Mantophryne | Callulops | 1 | 0 | 0 |
| Mantophryne | Mantophryne | 3 | 4 | 4 |
| Oreophryne | Asterophrys | 0 | 3 | 3 |
| Oreophryne | Choerophryne | 2 | 5 | 5 |
| Oreophryne | Cophixalus | 7 | 9 | 9 |
| Oreophryne | Oreophryne | 36 | 37 | 37 |
| Paedophryne | Paedophryne | 18 | 18 | 18 |
| Siamophryne | Cophixalus | 3 | 2 | 3 |
| Siamophryne | Oreophryne | 1 | 0 | 0 |
| Siamophryne | Siamophryne | 7 | 9 | 9 |
| Sphenophryne | Aphantophryne | 1 | 0 | 0 |
| Sphenophryne | Austrochaperina | 1 | 0 | 0 |
| Sphenophryne | Callulops | 1 | 0 | 0 |
| Sphenophryne | Choerophryne | 0 | 1 | 1 |
| Sphenophryne | Cophixalus | 1 | 0 | 0 |
| Sphenophryne | Copiula | 1 | 0 | 0 |
| Sphenophryne | Hylophorbus | 1 | 0 | 0 |
| Sphenophryne | Sphenophryne | 0 | 2 | 2 |
| Vietnamophryne | Austrochaperina | 1 | 1 | 1 |
| Vietnamophryne | Cophixalus | 2 | 3 | 3 |
| Xenorhina | Callulops | 1 | 0 | 0 |
| Xenorhina | Xenorhina | 24 | 25 | 25 |

**Figure S2.**
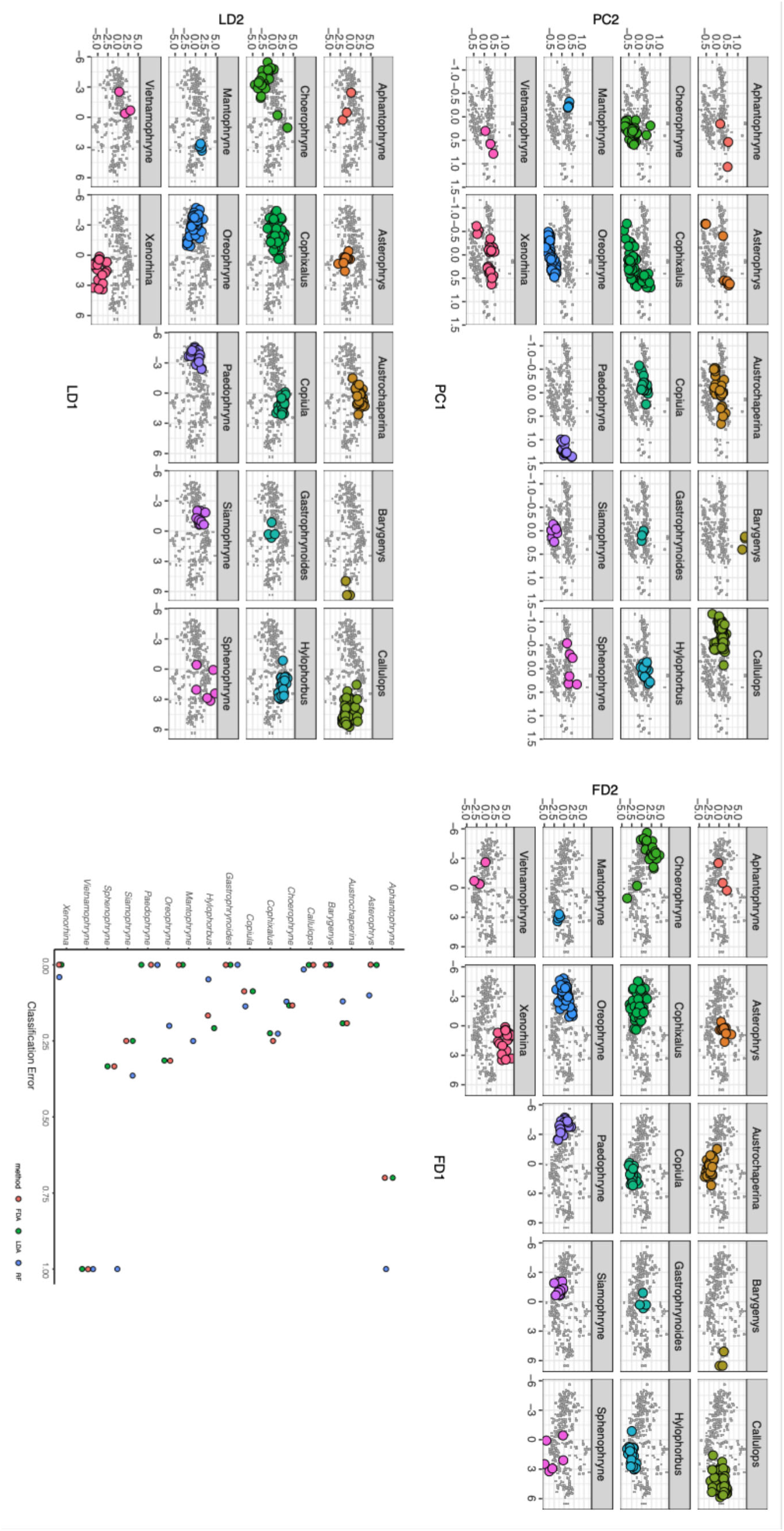
Comparison of visualization of dimensionality reduction techniques.

### Error and Bias

Measurement error and bias can be introduced to studies through a number of ways, includ-ing inter/intraobserver bias and lack of clarity around how measurements are taken. Haytek et al. (2001) highlights some of these sensitivities, particularly in inter/intraobserver bias. As a result, measurement error can go on to affect downstream interpretations. Our dataset comprises measurements taken across multiple decades by up to 13 different observers, leav-ing little ability to measure consistency, accuracy, and precision. Getting a single observer to re-measure dozens of individuals 20 or more times for hundreds of species is beyond the means of this study. In Haytek et al. (2001), the focus of the paper is sensitivity in intraspe-cific sampling. While we include intraspecific sampling in our dataset (none of which were originally published with supporting measurement error), our focus is above the species level. In this case the expectation is that interspecific variation (including error) is greater than intraspecific variation (including error), and that any error is random. To test this idea, we randomly downsampled our morphological dataset to 11 species with more than five indi-viduals measured per species (total 94 individuals). We used this dataset to run a separate Random Forest analysis where the classifier was the species. Of these 11 species, only one (Cophixalus balbus) had a classification error > 0. Looking closer, this is driven by a single individual which is not confidently assigned to the correct species (<0.70). Given that of 94 individuals across 11 species, only a single individual is misidentified, we can feel relatively confident that our assumption that interspecific (and inter-generic) variation is greater than intraspecific variation is true. We believe this provides evidence of the reliability of the data for our purpose.

**Figure S3.**
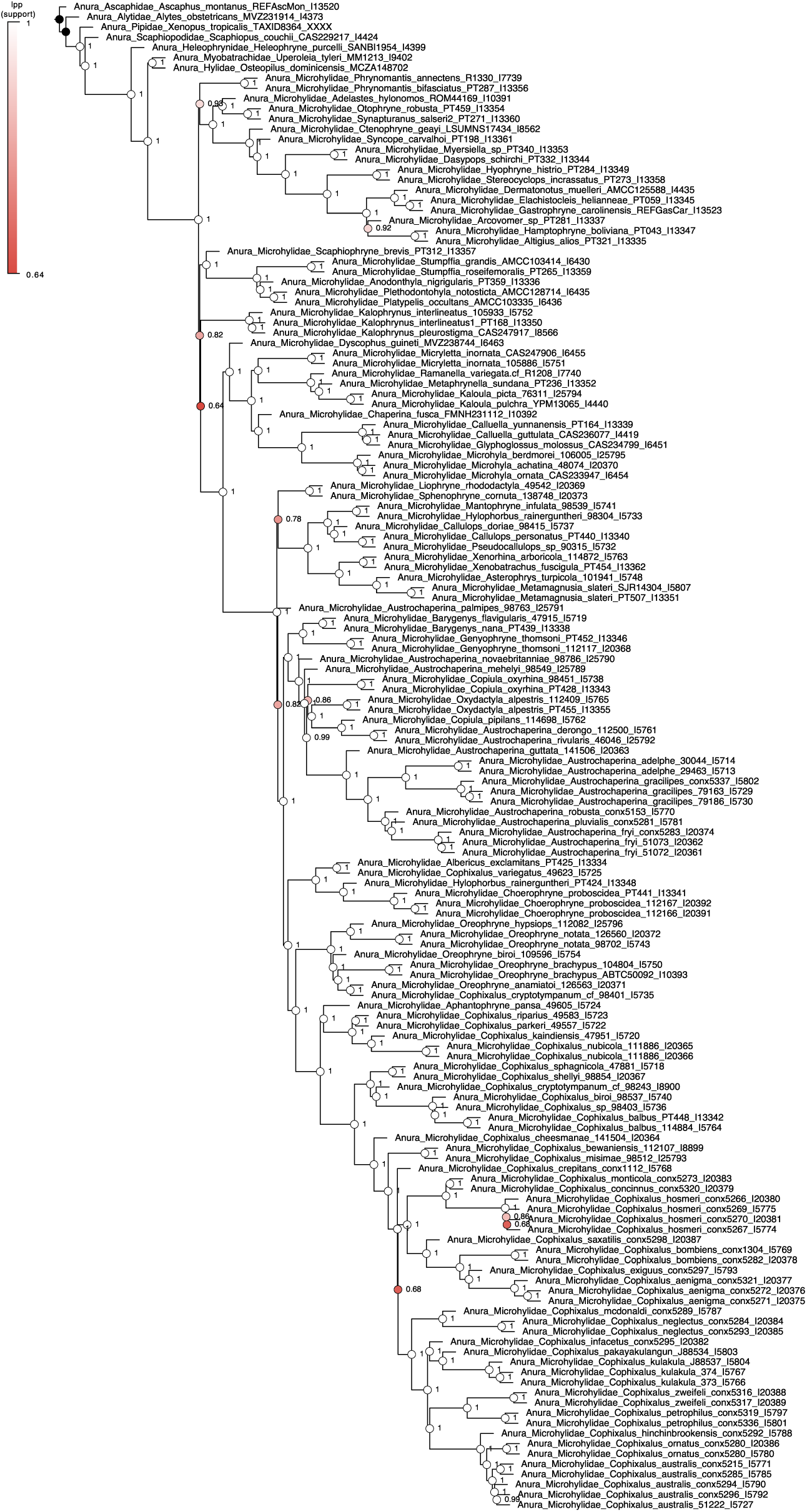
Fully sampled sequence capture tree, estimated by weighted hybrid ASTRAL using IQTREE genetree inputs. Branch support values are indicated at nodes and colored according to value (white = 1; red <= 0.9).

**Figure S4.**
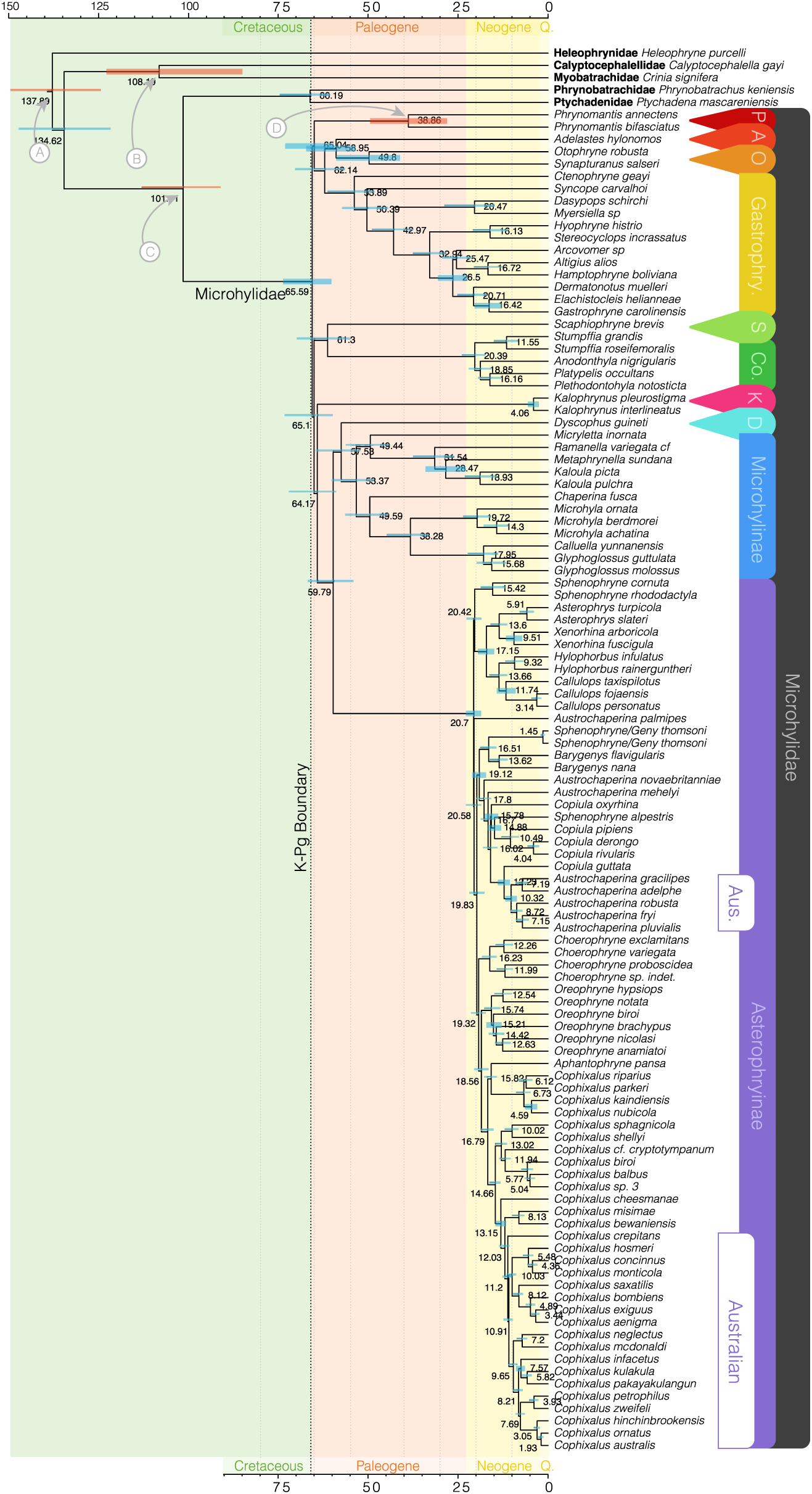
Time calibrated species tree of microhylid frogs highlights with confidence intervals indicated at nodes. Orange colored bars annotated with a circle indicate nodes calibrated by fossil evidence. These correspond to (A) *Beelzebufo ampinga* as a 66 million year minimum on the crown divergence of Neobatrachia; (B) *Calyptocephalella pichileufensis* as a 47.5 million year minimum on the split between Calyptocephalellidae and Myobatrachoidea; (C) *Thamatosaurus gezei* as a 33.9 million year minimum on the crown of Ranoidea; and (D) Ptychadenidae fossil as a 25 million year minimum on the split of Ptychadenidae and Phrynobatrachidae.

**Figure S5.**
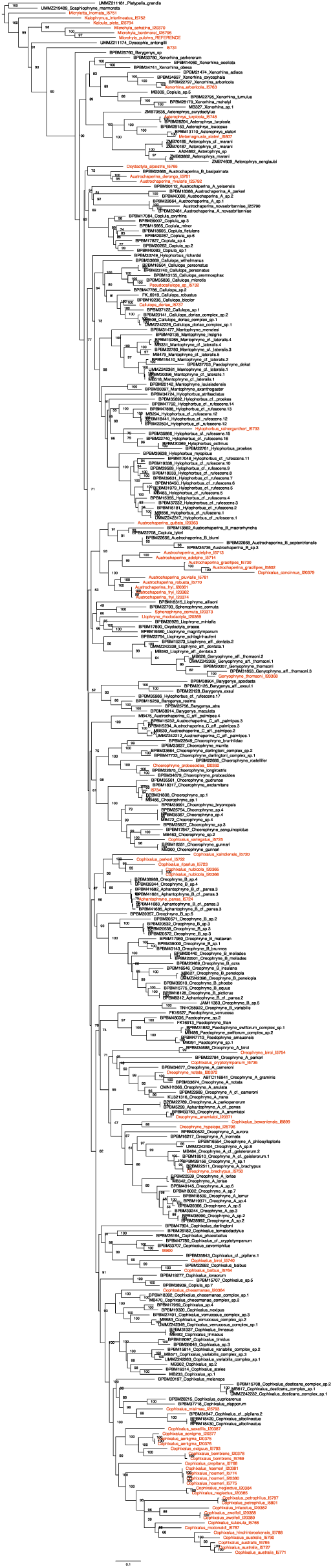
IQTREE gene tree of concatented mitochondrial loci (CYTB, ND4). Newly as-sembled and placed samples are indicated by orange text. Incorporating new samples into the alignments of Hill et al. (2023) allows for a shared understanding of Asterophryinae taxonomy between mitochondrial and nuclear datasets.

## Notes

https://github.com/iangbrennan/Asterophryinae

### Competing Interest Statement

The authors have declared no competing interest.

### Summary of Updates

Updates to figures, inclusion of additional references, and grammatical changes.

## References

1. Altenhoff, A.M., Warwick Vesztrocy, A., Bernard, C., Train, C.M., Nicheperovich, A., Prieto Baños, S., Julca, I., Moi, D., Nevers, Y., Majidian, S. and Dessimoz, C., (2024). OMA orthology in 2024: improved prokaryote coverage, ancestral and extant GO enrichment, a revamped synteny viewer and more in the OMA Ecosystem. Nucleic Acids Research, 52(D1), pp.D513–D521.

2. Anstis, M. (2017). Tadpoles and frogs of Australia. New Holland Publishers Pty Limited.

3. Benoit, M., Drost, H. G. (2021). A predictive approach to infer the activity and natural variation of retrotransposon families in plants. In: Cho J. (eds) Plant Transposable Elements. Methods in Molecular Biology, vol 2250. Humana, New York, NY.

4. Borowiec, M. L. (2016). AMAS: a fast tool for alignment manipulation and computing of summary statistics. PeerJ, 4, e1660.

5. Brennan, I. G., Lemmon, A. R., Moriarty Lemmon, E., Hoskin, C. J., Donnellan, S. C., & Keogh, J. S. (2024). Populating a continent: Phylogenomics reveal the timing of Australian frog diversification. Systematic Biology, 73(1), 1–11.

6. Burton, T. C. (1986). A reassessment of the Papuan subfamily Asterophryinae (Anura: Microhylidae). Records of the South Australian Museum, 19, 405–450.

7. Bushnell, B. (2014). BBMap: A Fast, Accurate, Splice-Aware Aligner. Berkeley, CA: Lawrence Berkeley National Lab.

8. Davies, H. L. (2012). The geology of New Guinea-the cordilleran margin of the Australian continent. Episodes Journal of International Geoscience, 35(1), 87–102.

9. de Sá, R. O., Streicher, J. W., Sekonyela, R., Forlani, M. C., Loader, S. P., Greenbaum, E., … & Haddad, C. F. (2012). Molecular phylogeny of microhylid frogs (Anura: Microhylidae) with emphasis on relationships among New World genera. BMC Evolutionary Biology, 12(1), 241.

10. dos Reis, M. & Yang, Z. (2011). Approximate likelihood calculation for Bayesian estimation of divergence times. Molecular Biology and Evolution, 28:2161–2172.

11. dos Reis, M., Gunnell, G. F., Barba-Montoya, J., Wilkins, A., Yang, Z., & Yoder, A. D. (2018). Using phylogenomic data to explore the effects of relaxed clocks and calibration strategies on divergence time estimation: primates as a test case. Systematic Biology, 67(4), 594–615.

12. Dubois, A., Ohler, A., & Pyron, R. A. (2021). New concepts and methods for phylogenetic taxonomy and nomenclature in zoology, exemplified by a new ranked cladonomy of recent amphibians (Lissamphibia). Megataxa, 5(1), 1–738.

13. Feng, Y. J., Blackburn, D. C., Liang, D., Hillis, D. M., Wake, D. B., Cannatella, D. C., & Zhang, P. (2017). Phylogenomics reveals rapid, simultaneous diversification of three major clades of Gondwanan frogs at the Cretaceous–Paleogene boundary. Proceedings of the National Academy of Sciences, 114(29), E5864–E5870.

14. Ferreira, F., Kraus, F., Richards, S., Oliver, P., Günther, R., Trilaksono, W., Arida, E.A., Hamidy, A., Riyanto, A., Tjaturadi, B. & Fouquet, A. (2024). Species delimitation and phylogenetic analyses of a New Guinean frog genus (Microhylidae: *Hylophorbus*) reveal many undescribed species and a complex diversification history driven by late Miocene events. Zoological Journal of the Linnean Society, 202(2), zlad168.

15. Ferreira, F., Oliver, P., Kraus, F., Günther, R., Richards, S., Tjaturadi, B., Arida, E., Hamidy, A., Riyanto, A., Trilaksono, W. and Thébaud, C., (2025). Molecular and acoustic evidence for large-scale underestimation of frog species diversity on New Guinea. Frontiers of Biogeography, 18, p.e137988.

16. Fonte, L. F. M. D., Mayer, M., & Lötters, S. (2019). Long-distance dispersal in amphibians. Frontiers of Biogeography, 11(4).

17. Frost, D.R. 2026. Amphibian Species of the World: an Online Reference. Version 6.2 (June 2026). Electronic Database accessible at https://amphibiansoftheworld.amnh.org/index.php. American Museum of Natural History, New York, USA. doi.org/10.5531/db.vz.0001

18. Gold, D. P., Casas-Gallego, M., Holm, R., Webb, M., & White, L. T. (2020). New tectonic reconstructions of New Guinea derived from biostratigraphy and geochronology. Proceedings of the Indonesian Petroleum Association Digital Technology Conference, 14-17 September 2020.

19. Günther, R., Dahl, C., & Richards, S. J. (2023). Another giant species of the microhylid frog genus *Cophixalus* Boettger, 1892 from the mountains of Papua New Guinea and first records of procoracoids in the genus. Zoosystematics and Evolution, 99(1), 173–183.

20. Hall, R. (1997). Cenozoic plate tectonic reconstructions of SE Asia. *Geological Society, London*, Special Publications, 126(1), 11–23.

21. Hayek, L. A. C., Heyer, W. R., & Gascon, C. (2001). Frog morphometrics: a cautionary tale. Alytes, 18(3-4), 153–177.

22. Hill, E.C., Fraser, C.J., Gao, D.F., Jarman, M.J., Henry, E.R., Iova, B., Allison, A. & Butler, M.A., (2022). Resolving the deep phylogeny: implications for early adaptive radiation, cryptic, and present-day ecological diversity of Papuan microhylid frogs. Molecular Phylogenetics and Evolution, 177, p.107618.

23. Hill, E. C., Gao, D. F., Polhemus, D. A., Fraser, C. J., Iova, B., Allison, A., & Butler, M. A. (2023). Testing geology with biology: plate tectonics and the diversification of microhylid frogs in the Papuan region. Integrative Organismal Biology, 5(1), obad028.

24. Hime, P. M., Lemmon, A. R., Lemmon, E. M., Prendini, E., Brown, J. M, Thomson, R. C, Kratovil, J. D, Noonan, B. P, Pyron, R A., Peloso, P. L V, Kortyna, M. L, Keogh, J. S., Donnellan, S. C, Mueller, R. L., Raxworthy, C. J, Kunte, K., Ron, S. R, Das, S., Gaitonde, N., Green, D. M, Labisko, J., Che, J., & Weisrock, D. W. (2021). Phylogenomics reveals ancient gene tree discordance in the amphibian tree of life. Systematic Biology, 70(1), 49–66.

25. Hoskin, C. J. (2004). Australian microhylid frogs (*Cophixalus* and *Austrochaperina*): phylogeny, taxonomy, calls, distributions and breeding biology. Australian Journal of Zoology, 52(3), 237–269.

26. Hoskin, C. J. (2013). A new frog species (Microhylidae: *Cophixalus*) from boulder-pile habitat of Cape Melville, north-east Australia. Zootaxa, 3722, 61–72.

27. IUCN. 2026. The IUCN Red List of Threatened Species. Version 2026-1. https://www.iucnredlist.org/resources/spatial-data-download.

28. Katoh, K., & Standley, D. M. (2013). MAFFT multiple sequence alignment software version 7: improvements in performance and usability. Molecular biology and evolution, 30(4), 772–780.

29. Kalyaanamoorthy, S., Minh, B. Q., Wong, T. K., Von Haeseler, A., & Jermiin, L. S. (2017). ModelFinder: fast model selection for accurate phylogenetic estimates. Nature Methods, 14(6), 587–589.

30. Köhler, F., & Günther, R. (2008). The radiation of microhylid frogs (Amphibia: Anura) on New Guinea: A mitochondrial phylogeny reveals parallel evolution of morphological and life history traits and disproves the current morphology-based classification. Molecular Phylogenetics and Evolution, 47(1), 353–365.

31. Kurabayashi, A., Matsui, M., Belabut, D. M., Yong, H. S., Ahmad, N., Sudin, A., … & Sumida, M. (2011). From Antarctica or Asia? New colonization scenario for Australian-New Guinean narrow mouth toads suggested from the findings on a mysterious genus *Gastrophrynoides*. BMC Evolutionary Biology, 11(1), 175.

32. Liaw, A., & Wiener, M. (2002). Classification and regression by randomForest. R news, 2(3), 18–22.

33. Linkem, C. W., Minin, V. N., & Leaché, A. D. (2016). Detecting the anomaly zone in species trees and evidence for a misleading signal in higher-level skink phylogeny (Squamata: Scincidae). Systematic Biology, 65(3), 465–477.

34. Lemmon, A. R., Emme, S. A., & Lemmon, E. M. (2012). Anchored hybrid enrichment for massively high-throughput phylogenomics. Systematic Biology, 61(5), 727–744.

35. Lewis, S.E., Sloss, C.R., Murray-Wallace, C.V., Woodroffe, C.D. & Smithers, S.G. (2013). Post-glacial sea-level changes around the Australian margin: a review. Quaternary Science Reviews, 74, 115–138.

36. Longrich, N. R., Bhullar, B. A. S., & Gauthier, J. A. (2012). Mass extinction of lizards and snakes at the Cretaceous–Paleogene boundary. Proceedings of the National Academy of Sciences, 109(52), 21396–21401.

37. Martin, H. A. (2006). Cenozoic climatic change and the development of the arid vegetation in Australia. Journal of Arid Environments, 66(3), 533–563.

38. Meredith, R. W., Janečka, J. E., Gatesy, J., Ryder, O. A., Fisher, C. A., Teeling, E. C., … & Murphy, W. J. (2011). Impacts of the Cretaceous Terrestrial Revolution and KPg extinction on mammal diversification. Science, 334(6055), 521–524.

39. Miller, K. G., Browning, J. V., Schmelz, W. J., Kopp, R. E., Mountain, G. S., & Wright, J. D. (2020). Cenozoic sea-level and cryospheric evolution from deep-sea geochemical and continental margin records. Science Advances, 6(20), eaaz1346.

40. Minh, B. Q., Nguyen, M. A. T., & von Haeseler, A. (2013). Ultrafast approximation for phylogenetic bootstrap. Molecular Biology and Evolution, 30(5), 1188–1195.

41. Minh, B. Q., Schmidt, H. A., Chernomor, O., Schrempf, D., Woodhams, M. D., Von Haeseler, A., & Lanfear, R. (2020). IQ-TREE 2: new models and efficient methods for phylogenetic inference in the genomic era. Molecular Biology and Evolution, 37(5), 1530–1534.

42. Mitchell, K.J., Pratt, R.C., Watson, L.N., Gibb, G.C., Llamas, B., Kasper, M., Edson, J., Hopwood, B., Male, D., Armstrong, K.N. and Meyer, M., (2014). Molecular phylogeny, biogeography, and habitat preference evolution of marsupials. Molecular Biology and Evolution, 31(9), pp.2322–2330.

43. Morris, R. S., Kraus, F., Deepak, V., Pal, S., Richards, S. J., & Maddock, S. T. (2026). Genetic Diversification in a New Guinean Frog Genus (*Mantophryne*, Microhylidae) was Driven by Ancient Tectonic Activity and Climate Reorganisation. Ecology and Evolution, 16(5), e73291.

44. Nguyen, L. T., Schmidt, H. A., Von Haeseler, A., & Minh, B. Q. (2015). IQ-TREE: a fast and effective stochastic algorithm for estimating maximum-likelihood phylogenies. Molecular Biology and Evolution, 32(1), 268–274.

45. Peloso, P. L., Frost, D. R., Richards, S. J., Rodrigues, M. T., Donnellan, S., Matsui, M., Raxworthy, C.J., Biju, S.D., Lemmon, E.M., Lemmon, A.R. & Wheeler, W. C. (2016). The impact of anchored phylogenomics and taxon sampling on phylogenetic inference in narrow-mouthed frogs (Anura, Microhylidae). Cladistics, 32(2), 113–140.

46. Portik, D. M., Streicher, J. W., & Wiens, J. J. (2023a). Frog phylogeny: a time-calibrated, species-level tree based on hundreds of loci and 5,242 species. Molecular Phylogenetics and Evolution, 188, 107907.

47. Portik, D. M., Streicher, J. W., Blackburn, D. C., Moen, D. S., Hutter, C. R., & Wiens, J. J. (2023b). Redefining possible: combining phylogenomic and supersparse data in frogs. Molecular Biology and Evolution, 40(5), msad109.

48. Poyarkov Jr, N. A., Suwannapoom, C., Pawangkhanant, P., Aksornneam, A., Van Duong, T., Korost, D. V., & Che, J. (2018). A new genus and three new species of miniaturized microhylid frogs from Indochina (Amphibia: Anura: Microhylidae: Asterophryinae). Zoological Research, 39(3), 130.

49. Prjibelski, A., Antipov, D., Meleshko, D., Lapidus, A., & Korobeynikov, A. (2020). Using SPAdes de novo assembler. Current Protocols in Bioinformatics, 70(1), e102.

50. Rannala, B., Yang, Z. (2007) Inferring speciation times under an episodic molecular clock. Systematic Biology, 56:453–466.

51. Ranwez, V., Douzery, E. J., Cambon, C., Chantret, N., & Delsuc, F. (2018). MACSE v2: toolkit for the alignment of coding sequences accounting for frameshifts and stop codons. Molecular Biology and Evolution, 35(10), 2582–2584.

52. Rivera, J. A., Kraus, F., Allison, A., & Butler, M. A. (2017). Molecular phylogenetics and dating of the problematic New Guinea microhylid frogs (Amphibia: Anura) reveals elevated speciation rates and need for taxonomic reclassification. Molecular Phylogenetics and Evolution, 112, 1–11.

53. Steenwyk, J. L., Buida III, T. J., Li, Y., Shen, X. X., & Rokas, A. (2020). ClipKIT: a multiple sequence alignment trimming software for accurate phylogenomic inference. PLoS biology, 18(12), e3001007.

54. Streicher, J. W., Loader, S. P., Varela-Jaramillo, A., Montoya, P., & de Sá, R. O. (2020). Analysis of ultraconserved elements supports African origins of narrow-mouthed frogs. Molecular Phylogenetics and Evolution, 146, 106771.

55. Suwannapoom, C., Sumontha, M., Tunprasert, J., Ruangsuwan, T., Pawangkhanant, P., Korost, D. V., & Poyarkov, N. A. (2018). A striking new genus and species of cave-dwelling frog (Amphibia: Anura: Microhylidae: Asterophryinae) from Thailand. PeerJ, 6, e4422.

56. Tu, N., Yang, M., Liang, D., & Zhang, P. (2018). A large-scale phylogeny of Microhylidae inferred from a combined dataset of 121 genes and 427 taxa. Molecular Phylogenetics and Evolution, 126, 85–91.

57. Tyler, M. J. (1998). Australian Frogs: A Natural History. Cornell University Press.

58. van der Meijden, A., Vences, M., Hoegg, S., Boistel, R., Channing, A., & Meyer, A. (2007). Nuclear gene phylogeny of narrow-mouthed toads (Family: Microhylidae) and a discussion of competing hypotheses concerning their biogeographical origins. Molecular Phylogenetics and Evolution, 44(3), 1017–1030.

59. VanDerWal, J., Shoo, L. P., & Williams, S. E. (2009). New approaches to understanding late Quaternary climate fluctuations and refugial dynamics in Australian wet tropical rain forests. Journal of Biogeography, 36(2), 291–301.

60. Wallace, A. R. (1869). The Malay Archipelago: The land of the orang-utan, and the bird of paradise. A narrative of travel, with studies of man and nature. Macmillan.

61. Wickham, H. (2011). ggplot2. Wiley interdisciplinary reviews: computational statistics, 3(2), 180–185.

62. Wu, J., Yonezawa, T., & Kishino, H. (2017). Rates of molecular evolution suggest natural history of life history traits and a post-K-Pg nocturnal bottleneck of placentals. Current Biology, 27(19), 3025–3033.

63. Zhang, C., Rabiee, M., Sayyari, E., & Mirarab, S. (2018). ASTRAL-III: polynomial time species tree reconstruction from partially resolved gene trees. BMC Bioinformatics, 19(6), 15–30.

64. Zweifel, R. G. (1972). A revision of the frogs of the subfamily Asterophryinae, family Microhylidae. Bulletin of the American Museum of Natural History; v. 148, article 3.

65. Zweifel, R. G. (1985). Australian frogs of the family Microhylidae. Bulletin of the American Museum of Natural History; v. 182, article 3.

66. Zweifel, R. G. (2000). Partition of the Australopapuan microhylid frog genus *Sphenophryne* with descriptions of new species. Bulletin of the American Museum of Natural History, 2000(253), 1–130.

